# A novel platform for metabolomics using barcoded structure-switching aptamers

**DOI:** 10.1101/2023.06.09.544402

**Authors:** June H. Tan, Maria P. Mercado, Andrew G. Fraser

## Abstract

Small organic molecules like metabolites and drugs are critical for diagnostics, treatment, and synthetic biology. Measuring them presents two key challenges however: they are biochemically highly diverse and there is no method to amplify them. Mass spectrometry has been the workhorse of metabolomics for decades but is costly and slow and single-cell metabolomics remains very challenging. Here we describe an alternative platform for metabolomics based on structure-switching aptamers (SSAs). SSAs are short nucleic acid molecules that each recognise a specific target ligand and undergo a major conformational change on ligand binding. This conformational change can drive detection such as fluorescence allowing SSAs to be used as sensors. We adapted conventional SSAs to a novel readout: barcode release. Each SSA recognises a unique ligand and each SSA releases a unique barcode allowing many ligands to be detected in parallel. We show that these barcode SSAs (bSSAs) can be multiplexed and act as independent sensors and that barcode release can be massively amplified to allow high sensitivity. Finally, we establish methods for the generation of large collections of bSSAs where barcode-SSA matching is completely directed. We believe that this novel platform which converts metabolite detection into barcode sequencing will allow the deep multiplexed detection of metabolites and drugs down to the scale of single cells.

## Introduction

Small organic molecules like metabolites and drugs are at the heart of biology, and there is an urgently growing need for effective ways to measure their levels down to the scale of single cells^1, 2^. The standard workhorse for detecting these small molecules is mass spectrometry (MS)^2–4^. MS is extremely powerful but has limitations that are intrinsic to the chemistry of metabolites and drugs. In particular, MS has limits to sensitivity which makes detection of low levels of these molecules very challenging. This is a key issue for measuring small molecules in single cells^1, 5, 6^. Single cell biology has exploded as a field in the last decade and being able to measure mRNA levels and sequence genomes in single cells has given key insights into tumorigenesis and developmental biology^7–10^. However, while it is easy to analyse DNA and mRNAs in single cells, it is a much greater challenge for small molecules. The reason is simple: it is trivial to amplify nucleic acids using PCR, but there is no equivalent ‘PCR for metabolites’. To make single-cell metabolomics as simple and sensitive as single-cell RNA-seq (scRNA-seq), we need some simple universal way to amplify the signal of small molecules.

Metabolites and drugs are biochemically very diverse — it is impossible to imagine a universal way to directly amplify sugars, amino acids, lipids, and every different drug chemistry in the same way that PCR can amplify any DNA sequence. We have therefore developed a different approach. Conceptually, if we could read out the complex biochemistry of metabolites and drugs with DNA barcodes, this would allow us to PCR amplify the signal. This type of approach is already used for drug screening with ‘DNA-encoded’ drug libraries^11–14^. Each drug is coupled to a unique known DNA barcode and thus it is possible to detect which drugs bound to a target protein by simply reading out which barcodes are present^13, 15^. That has proved powerful but there is a critical difference with metabolites: metabolites are not intrinsically barcoded. In the DNA-encoded drug libraries, each drug is made with a barcode attached; in a cell lysate or blood sample, for example, the metabolites have no barcode. If we could somehow pick up every glucose molecule in a sample and add a unique barcode to them, then pick up every phenylalanine molecule and add a different barcode, we could read out their levels using DNA barcodes. How can we recognise every metabolite in a sample and couple it to a unique DNA barcode? We report here a universal method for coupling metabolites and drugs to barcodes using barcoded Structure Switching Aptamers (bSSAs).

Aptamers are short oligonucleotides, typically made of DNA, or RNA, that bind specific targets like metabolites or drugs with high affinity^16, 17^. Structure-switching aptamers are unique in that they undergo a very large conformational change when their target ligand is bound by the ligand-binding region of the aptamer. This conformational change can be detected and used as a readout of ligand binding — typically this is fluorescence, but other common readouts include conductance and other biophysical changes^18, 19^. SSAs are thus powerful tools for sensing small molecules and drugs and SSAs have been developed as sensors for phenylalanine^20^, theophylline^21, 22^ and kanamycin^23^ for example.

Previous SSA readouts have two key limitations, however. First, every SSA has the same readout — it is impossible to multiplex SSAs if they all have the same fluorescent signal, and since there are thousands of different metabolites in a single cell, we need sensors with non-overlapping readouts if we want to be able to see them all in parallel. Second, while many SSAs are very sensitive, their signal cannot be trivially amplified. This is a specific problem for single cell metabolomics, where the amount of the starting ligand is very limited^24^. Here, we report a new readout for ligand-binding to SSAs — barcode release. Each target ligand is recognised by an SSA that contains a unique barcode. Ligand binding causes the release of the DNA barcode – by sequencing the barcodes that are released upon ligand binding, we can infer the levels of the associated small molecules in the sequenced sample. These barcodes can be easily amplified, and many thousands of sensors can be analysed in parallel since each releases a different barcode. We show that barcode release provides an excellent way to quantify ligand levels, that bSSAs can be multiplexed and read out in parallel, and provide methods for the bulk assembly of bSSAs that correctly pair barcodes with ligand-binding regions. We believe this approach has the potential to transform single-cell metabolomics and since the readout is DNA barcode release, metabolites can be measured and read on the same platform as scRNA-seq.

## Results

### Barcode release is a novel readout for structure-switching aptamers

Structure-switching aptamers undergo a major conformational change following ligand binding. Detecting this conformational change allows them to be used as sensors^20, 25, 26^. One method to detect the conformational change in an SSA due to ligand binding is to sense the dissociation of a bimolecular SSA into its two separate parts — the ligand binding oligo (LBO from here on) and a short release oligo (SRO from here on) (Fig 1). The LBO provides the ligand-binding region that confers ligand specificity and the SRO is coupled to a universal readout. The LBO and SRO interact via base-pairing, which is then disrupted by the conformational change following ligand binding. That dissociation can be detected using various methods, many of which commonly involve fluorescence (Fig 1A), but spectroscopic and electrical conductance methods are also frequently used^20, 27, 28^. One difficulty with all these detection methods is that it is not possible to assay many different SSAs in a single physical location (e.g., a single sample tube) if all the SSAs have the same readout. We solve this by introducing a new way to read out the binding of a ligand to an SSA — DNA barcode release.

**Fig 1.**
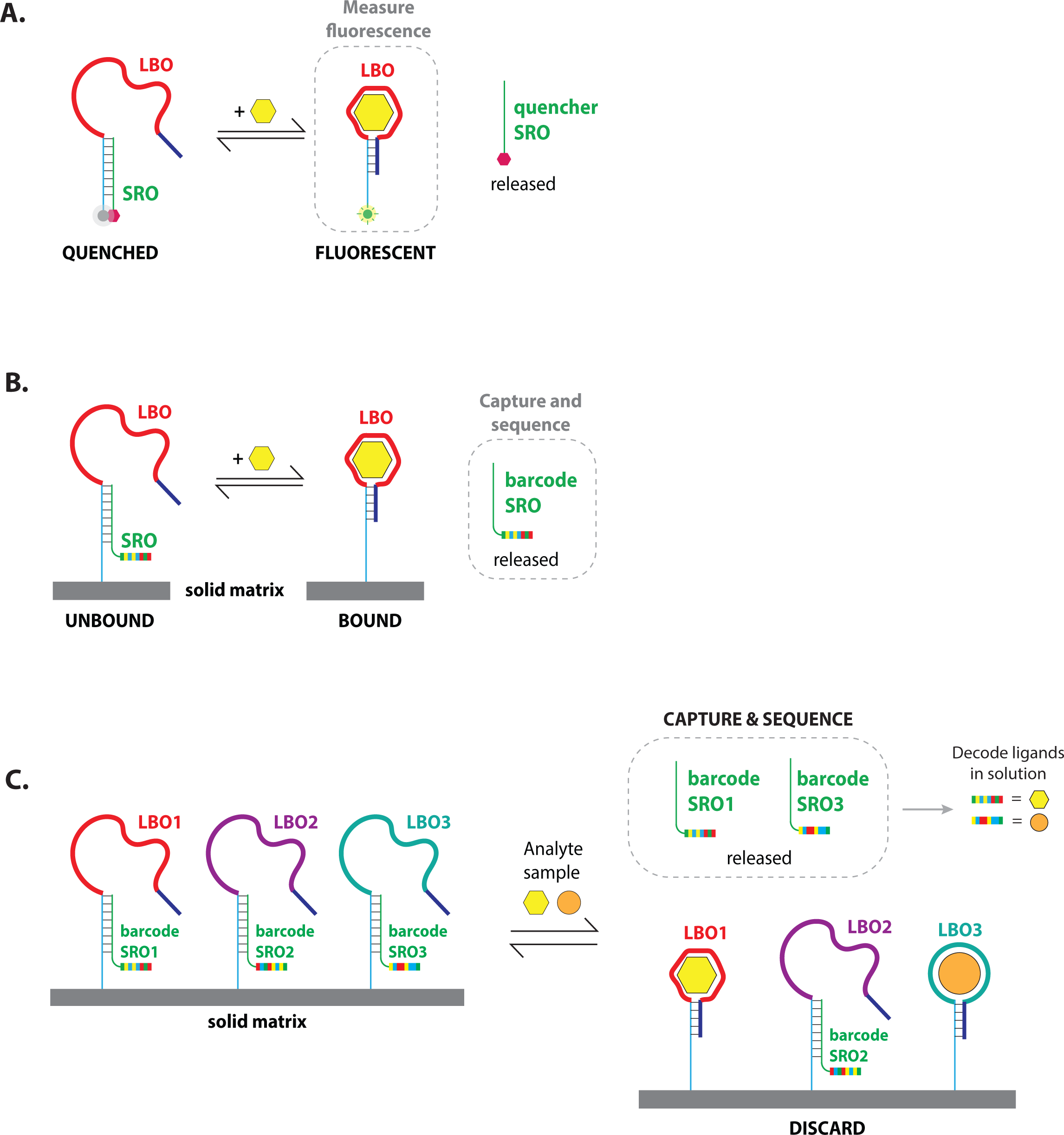
Architecture of barcoded structure-switching aptamers (bSSAs) Each structure-switching aptamer (SSA) is comprised of two parts – a ligand binding oligo (LBO) and a short-release oligo (SRO) which bind via base-pairing. When the LBO is bound by its cognate ligand (yellow hexagon), a conformation change induces the release of an SRO. **(A) Ligand binding is monitored in solution using a fluorophore-quencher pair.** The LBO is attached to a fluorophore that is quenched when base-paired with an SRO with a quencher. Ligand binding to the LBO induces the displacement of the SRO quencher and results in an increase in fluorescence signal. **(B) Ligand binding is monitored using an SRO with a unique barcode sequence**. The barcode SRO is base-paired to the LBO that is attached to a solid matrix. Here, the LBO is biotinylated at its 5′ end and attached to streptavidin-coated magnetic beads. Ligand binding displaces the barcode SRO that contains a sequence specific to that sensor, which can then be amplified and sequenced. **(C) bSSAs can be multiplexed and used to detect ligand binding in parallel for a large library of SSAs.** Ligand binding to the LBO results in release of the corresponding barcode SRO, which can then be amplified and sequenced. Sequencing of released barcodes would show which SSAs bound their ligand and consequently identify ligands that were present in the sample.

Instead of using a universal SRO, each SSA has a different SRO. These all have the same base-pairing region to bind the LBO but have an additional unique barcode (Fig 1B). This allows multiplexing e.g., if a glucose-binding LBO is attached to an SRO with one barcode and a galactose-binding LBO is bound to an SRO with a different barcode, we can read out glucose and galactose levels in parallel simply by detecting and sequencing the released barcoded SROs (schematic in Fig 1C). To make this work in practice, libraries of bimolecular bSSA molecules are all attached to a solid matrix, such as to streptavidin-coated beads via a 5′ biotinylated end. Ligand-binding drives release of barcode SROs and these can then be separated and sequenced. Since any specific bSSA is present in many thousands of identical copies, this provides a digital readout of ligand levels: the more released barcode we see, the more molecules of ligand bound their sensor. This provides ample dynamic range to measure thousands of metabolites in parallel — for example it is possible to have >10 million sensor molecules on a single bead or well of a microfluidics device^29^. Finally, released barcodes are entirely comprised of DNA and can be readily amplified by PCR — this has potential for single cell metabolomics.

To test whether our bSSA architecture works, we first established a fluorescence-based assay to detect released SRO in lieu of sequencing the released barcodes. This allows for a more rapid and cost-effective way to measure SRO release for simple proof-of-concept experiments. We thus established the method shown in Fig 2. Instead of using an arbitrary sequence barcode attached to the SRO, we use a ‘barcode’ that corresponds to the sequence of the T7 RNA polymerase promoter^30^ (‘T7 SRO’ from here on) (Fig 2A). To detect release, we use a reporter containing the anti-sense T7 promoter region and a transcription template that directs the transcription of the Baby Spinach^31^ RNA aptamer. When the T7 SRO is released, it can anneal to this template — the region of the T7 promoter is now double-stranded on base-pairing to the T7 SRO, allowing it to act as a promoter for transcription^32^ to drive transcription of the Baby Spinach RNA aptamer in an in vitro transcription reaction (Fig 2B). Baby Spinach^31, 33^, a derivative of the Spinach aptamer^34^, is a 51 nt fluorogenic RNA that fluoresces when complexed with the fluorophore DFHBI or its derivatives (e.g. DFHBI-1T). In this way, SSA dissociation can be monitored simply by fluorescence: T7 SRO release leads to Baby Spinach transcription which is detected by fluorescence. As shown in Fig 2C, titration of the T7 SRO into a reaction containing excess Baby Spinach reporter results in dose-dependent fluorescence, and thus, this method allows us to monitor SSA ligand binding efficiently by estimating levels of T7 SRO released.

**Fig 2.**
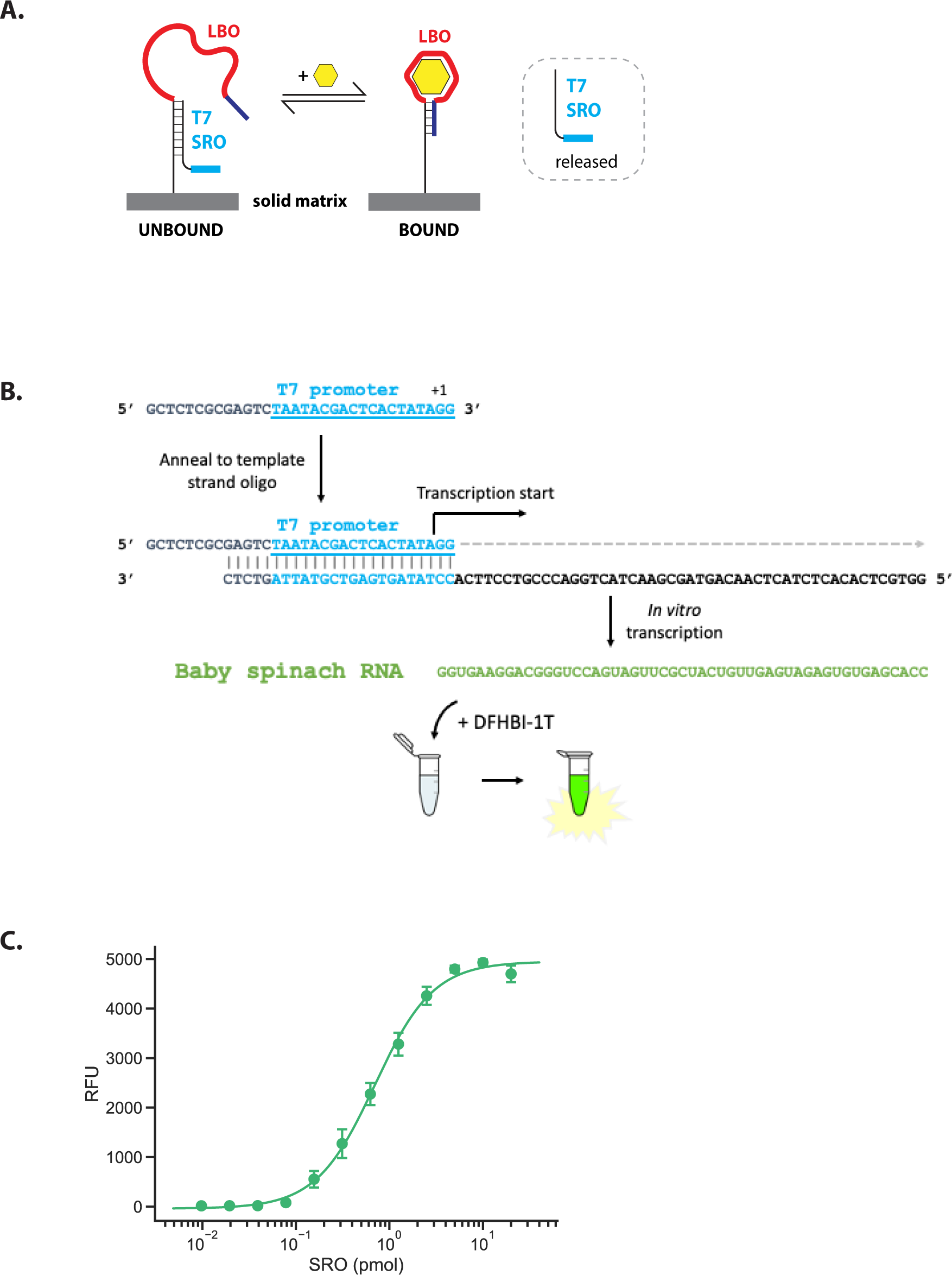
Readout of barcode SRO release using a fluorescent reporter. **(A) Schematic of SSAs used in proof-of-concept experiments.** To detect amounts of released barcode SRO, we used a T7 promoter sequence in place of the barcode sequence. This T7 promoter sequence can then be added to a transcription template to allow transcription of any reporter sequence. **(B) Schematic of T7-Baby Spinach fluorescence assay.** To detect SRO release using a fluorescent reporter, the SRO sequence containing a T7 promoter is annealed to a transcription template for Baby Spinach. The Baby Spinach RNA is then transcribed by T7 RNA polymerase in an *in vitro* transcription reaction. DFHBI-1T fluoresces only when bound to the folded Baby Spinach aptamer, allowing detection of the transcribed products. **(C) Increased fluorescence is observed with increasing amounts of T7 SRO input.** A series of 2-fold dilutions of T7 barcode SROs added to 20 pmol of Baby Spinach transcription template. After a 2h *in vitro* transcription reaction, the transcribed products were heated at 90°C for 2 min, then placed on ice for 5 min. DFHBI-1T was then added at a final concentration of 10 µM, and the mixture was then heated at 65°C for 5 min and slowly cooled to room temperature. Fluorescence intensity for each sample was then measured using a FLUOstar Omega microplate reader (ex 485nm, em 520nm). Error bars show standard error.

To assess whether bSSAs can be used to detect a range of different ligands, we first adapted published aptamers that bind either endogenous metabolites (e.g. phenylalanine^20^, cortisol^35^, ATP^36^) or exogenous targets (e.g., the antibiotic ampicillin^37^ and the antimalarial drugs piperaquine^38^, and mefloquine^38^) onto our bSSA platform. Using the T7-Spinach readout shown in Fig 2B, we show that barcode release gives a quantitative readout of ligand levels in solution (Fig 3A). To confirm that this method does not skew levels of SRO abundance due to any non-linear amplification, we compared the dose response curve to sensors with a direct, non-amplified fluorescence output. We find that our T7-Spinach assay results in a very similar dose response to a direct fluorescence readout, suggesting that our linear amplification method should not greatly skew levels of barcode release (Fig 3B).

**Fig 3.**
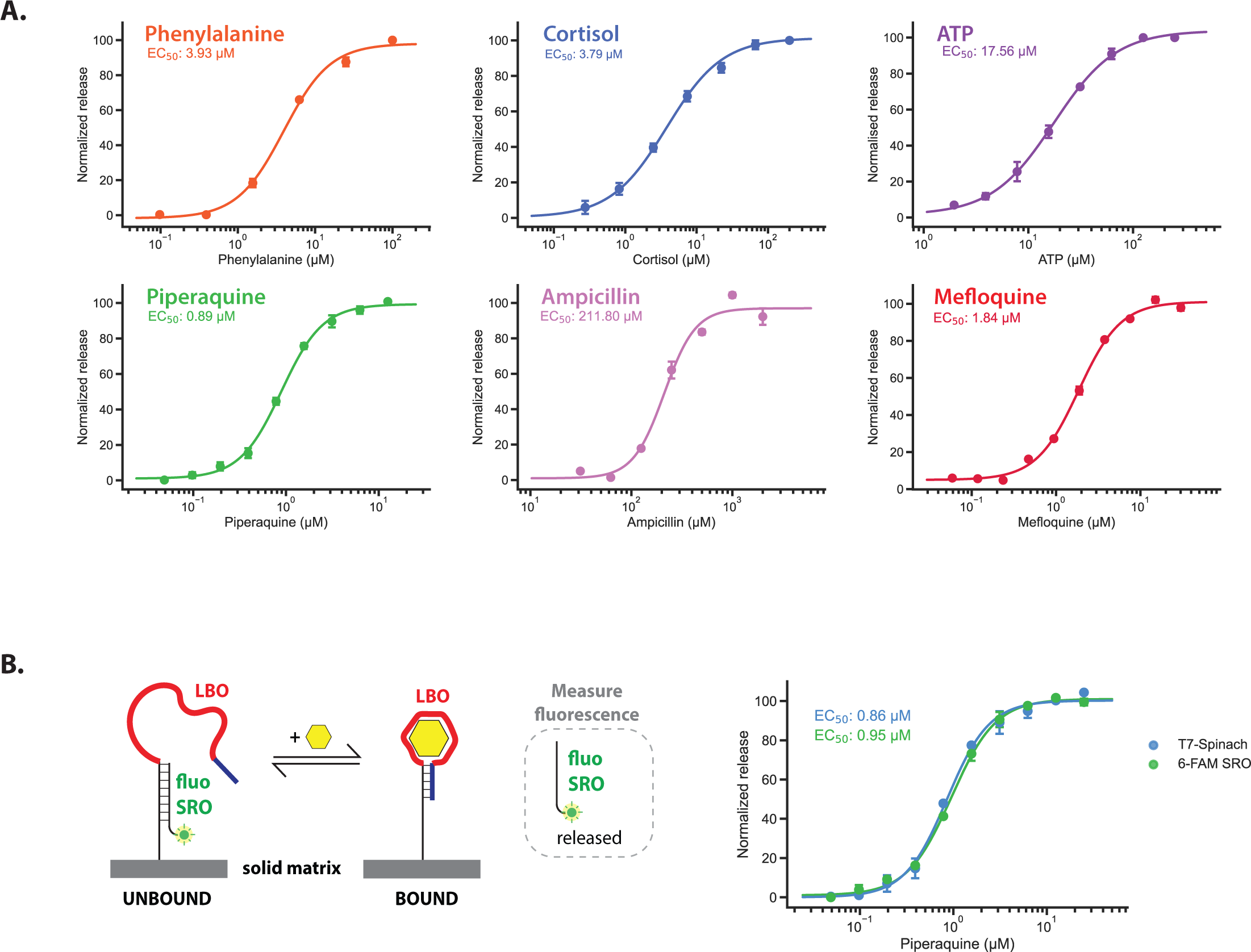
(A) T7 barcode SRO is released specifically in response to ligand. The ligand-binding region of LBOs were replaced by sequences from published aptamer sequences (see Methods and Supplemental File 1 for details). Dose response curves of each bSSAs to their cognate ligands were then measured using the assay described in Fig 2B, except for the cortisol and ATP DRCs, which were generated using a direct fluorescence readout described in Fig 3B. Relative proportions of SRO release were plotted by comparing the amount of SRO released in the supernatant with the amount of SRO that remained bound to beads (detailed in Methods). A four-parameter logistic (4PL) model was used for curve fitting and for estimating EC_50_ values. **(B) Comparison a direct fluorescent readout.** We compared the response of a piperaquine bSSA with a T7 readout with the same bSSA with an SRO attached to a 6-FAM fluorophore. For the latter, release of SRO was directly measured using a microplate reader (ex 485nm, em 520nm). Error bars in (A-B) show standard error from at least 3 independent replicates.

Additionally, we show that these bSSAs are highly specific for their target ligands, recapitulating the characterised specificity of their adapted aptamers^20, 37^ (Fig 4). These bSSAs can specifically distinguish their intended ligand from similarly structured compounds - the phenylalanine (Phe) bSSA can distinguish Phe from other aromatic amino acids like tyrosine (Tyr) (Fig 4A) and even between the L- and D-form enantiomers (Fig 4B) while the ampicillin (Amp) bSSA can distinguish its target from other close analogues that differ only in a single chain group (Fig 4C). We conclude that barcode release provides a faithful and accurate readout for ligand binding to a structure-switching aptamer and thus barcode release can be used to measure ligand levels.

**Fig 4.**
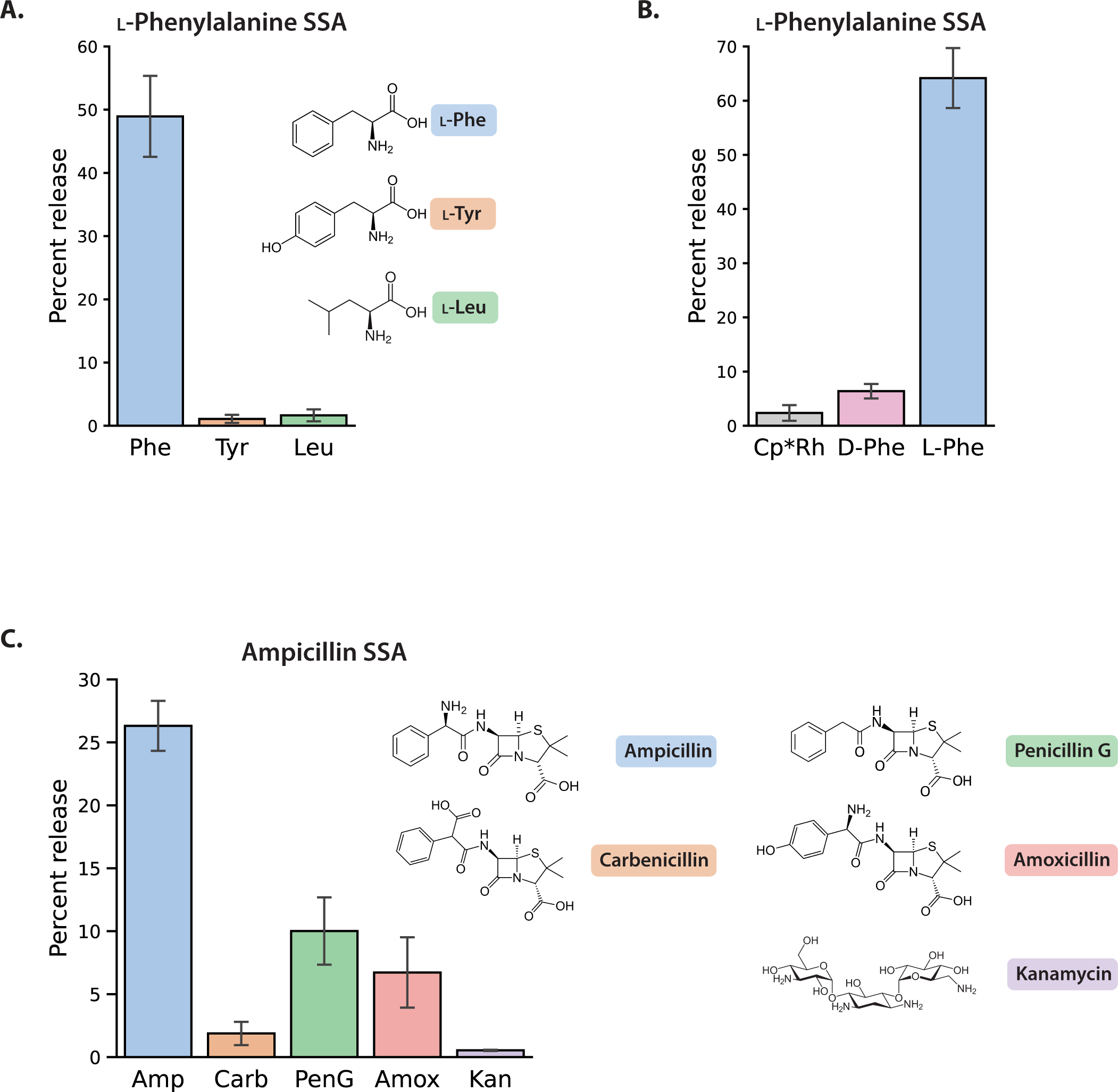
Specificity of bSSAs adapted from known aptamers. **(A) Specificity of the L-phenylalanine (Phe) bSSAs in response to various amino acids.** 10 µM of the respective amino acids were first complexed with 100 µM Cp*Rh(III) as described in Yang *et al.* 2014^20^, before incubation with the bound Phe bSSAs. The supernatant was collected after a 45 min incubation with ligand (supernatant fraction). The beads were then resuspended and boiled for 4 min to release all remaining SRO (remainder fraction). Percent release represents the fluorescence intensity detected in the supernatant fraction relative to the fluorescence intensity detected in both the supernatant fraction and the remaining bound fraction. Specific SRO release is observed in response to 10 µM Phe and not L-tyrosine (Tyr) or L-leucine (Leu) addition. **(B) Specificity of the L-Phe bSSAs in response to L- and D-enantiomers.** L-Phe bSSA was incubated with 100 µM of L-Phe or D-Phe complexed with 100 µM Cp*Rh(III) and assayed as described in (A). **(C) Specificity of the ampicillin (Amp) bSSA in response to various penicillin analogues.** The Amp bSSA was incubated with 500 µM of Amp, carbenicillin (Carb), penicillin G (PenG) or amoxicillin (Amox) or kanamycin (Kan) and assayed as described above. Error bars in (A-C) represent standard error from 3 independent replicates.

### Converting a unimolecular pre-SSA to a mature SSA results in an active SSA with a barcode readout

DNA barcodes have been used as readouts in a wide variety of experiments from large-scale RNAi screens to pooled yeast deletion screens to drug screens. The entire DNA barcoding approach relies on knowing the pairing between DNA barcode and the entity being measured e.g., drug X has barcode 1, drug Y has barcode 2 or yeast deletion A has barcode 3 and yeast deletion B has barcode 4. bSSAs are no different — the approach only works if we can correctly pair each barcode SRO with the ligand-binding LBO. This is simple to do for a single bSSA by mixing one barcoded SRO with one LBO with known ligand specificity and hybridising them together. However, this would be laborious if we want to make libraries of thousands of sensors with known barcode SRO-ligand-binding LBO pairings.

One possible solution is to have a unique base-pairing sequence between each barcode SRO and its LBO. A pool of SROs and LBOs could thus be mixed and in theory it might be possible for each SRO to hybridize uniquely to its correct LBO partner. However, the sequence space for the hybridising region is highly restricted since these different base-pairing regions must all have very similar sequence composition to ensure similar hybridisation and release properties; secondary structures and other constraints further limit the possible sequences. In addition, base pairing of each SRO to its matched LBO has to be error-free. If SROs frequently associate with incorrect LBOs, then it will be challenging to deconvolute since there is no longer a known match of barcode SRO to LBO and hence no longer a known match between barcode and bound ligand.

Here we take a different approach to ensure accurate pairing of LBO and an SRO with known barcode. Instead of synthesising LBO and barcode SRO as separate molecules and pairing them after synthesis, we make the bimolecular bSSAs as unimolecular pre-bSSAs — the barcode SRO and ligand-sensing LBO are thus intrinsically paired (Fig 5A). We fold the pre-SSAs and process them into the mature bimolecular bSSA by cleaving a single strand of the pre-bSSA either at an abasic site or using a nicking restriction enzyme. In this way, we can make large libraries of bimolecular bSSAs with known pairing between ligand-sensing LBOs and barcode SROs (bSROs). We compared the response of a bSSA that was assembled using separate LBO and barcode bSRO oligonucleotides with the response of a bSSA that was generated from a unimolecular pre-bSSA oligonucleotide. First, we show that the tryptophan (Trp) bSSA works when constructed from separate LBO and bSRO parts (Fig 5B). As control, we also generated a bSSA with scrambled version of the Trp ligand-binding region and observed negligible barcode release in response to Trp. We then synthesized the same Trp bSSA sequence but as a unimolecular starting molecule with an abasic site, processed it to its mature form using Endonuclease IV, and tested it in our assay. We find the resulting bSSA responds to Trp very similarly as the version made by hybridising the individual LBO and SRO oligos (Fig 5C). We therefore conclude that we can make mature functional SSAs with known SRO-LBO pairings by taking unimolecular pre-SSAs and converting them to mature SSAs. This can be done for libraries of any arbitrary size and allows us to efficiently produce large collections of SSAs where ligand binding can be monitored by assessing the release of barcoded SROs.

**Fig 5.**
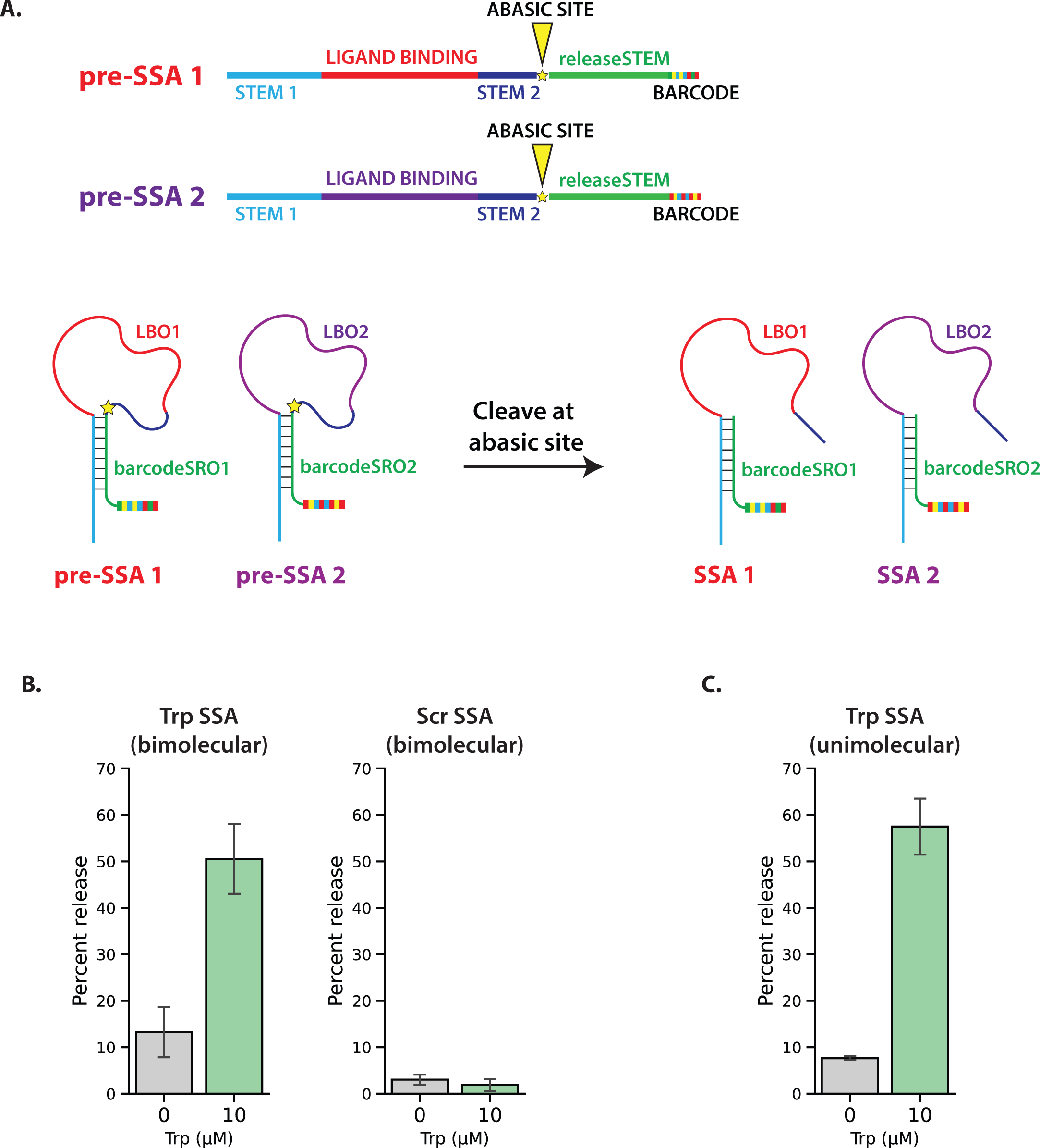
Assembly of bSSAs by cleaving a unimolecular pre-SSA to generate functional, paired LBO and SRO sequences. **(A) Schematic of unimolecular pre-bSSAs.** A library of barcoded SSAs can be generated by first synthesizing a pre-bSSA oligonucleotide that contains both the LBO and SRO sequences. These pre-SSA sequences each contain an abasic site (or other cleavage sites) that can be cleaved to generate the bimolecular mature bSSA construct. These pre-SSAs are then folded so that the LBO and SRO sequences are base paired. The folded constructs are then cleaved (e.g., at an abasic site using Endonuclease IV), thereby generating mature bSSAs. **(B) Trp-mediated SRO release from Trp bSSAs assembled from separate LBO and SRO oligonucleotides.** (Left) SRO release from Trp bSSAs was observed upon addition of 10 μM Trp complexed with Cp*Rh(III). (Right) Minimal Trp-mediated SRO release was observed from bSSAs with a randomized ligand-binding domain. **(C) Trp-mediated SRO release from Trp bSSAs processed and assembled from unimolecular pre-bSSAs.** The pre-SSA assembly method depicted in (A) was used to generate the same resulting Trp bSSA sequence as used in (B). With 10 μM Trp-Cp*Rh(III) addition, the mature Trp bSSA shows SRO release at a very similar level as constructs assembled from separate bimolecular parts. Error bars represent standard error from 3 independent replicates.

### Ligand detection in complex samples

In Fig 3, we show that barcode release from bSSAs can accurately read out ligand levels in a pure buffered solution. To be useful as sensors these bSSAs must also work in complex samples like cell lysate that contain thousands of other molecules. To test this, we titrated in piperaquine (PQ) in either a cell lysate mixture or LB broth solution and measured the response of the PQ bSSA (Fig 6A). After signal normalization, we observe very similar dose response curves in buffer, cell lysate and LB, suggesting that bSSAs work well even in the context of cell lysate or other complex medium. As further proof-of-concept, we treated *C. elegans* worms with PQ or a no-drug control to see if we could detect drug uptake in crude worm lysate using our PQ bSSA (Fig 6B). We observe much higher levels of barcode release in the lysate of worms treated with PQ relative to the no-drug control, further showing that our sensors can work in complex biological samples to detect their cognate ligand.

**Fig 6.**
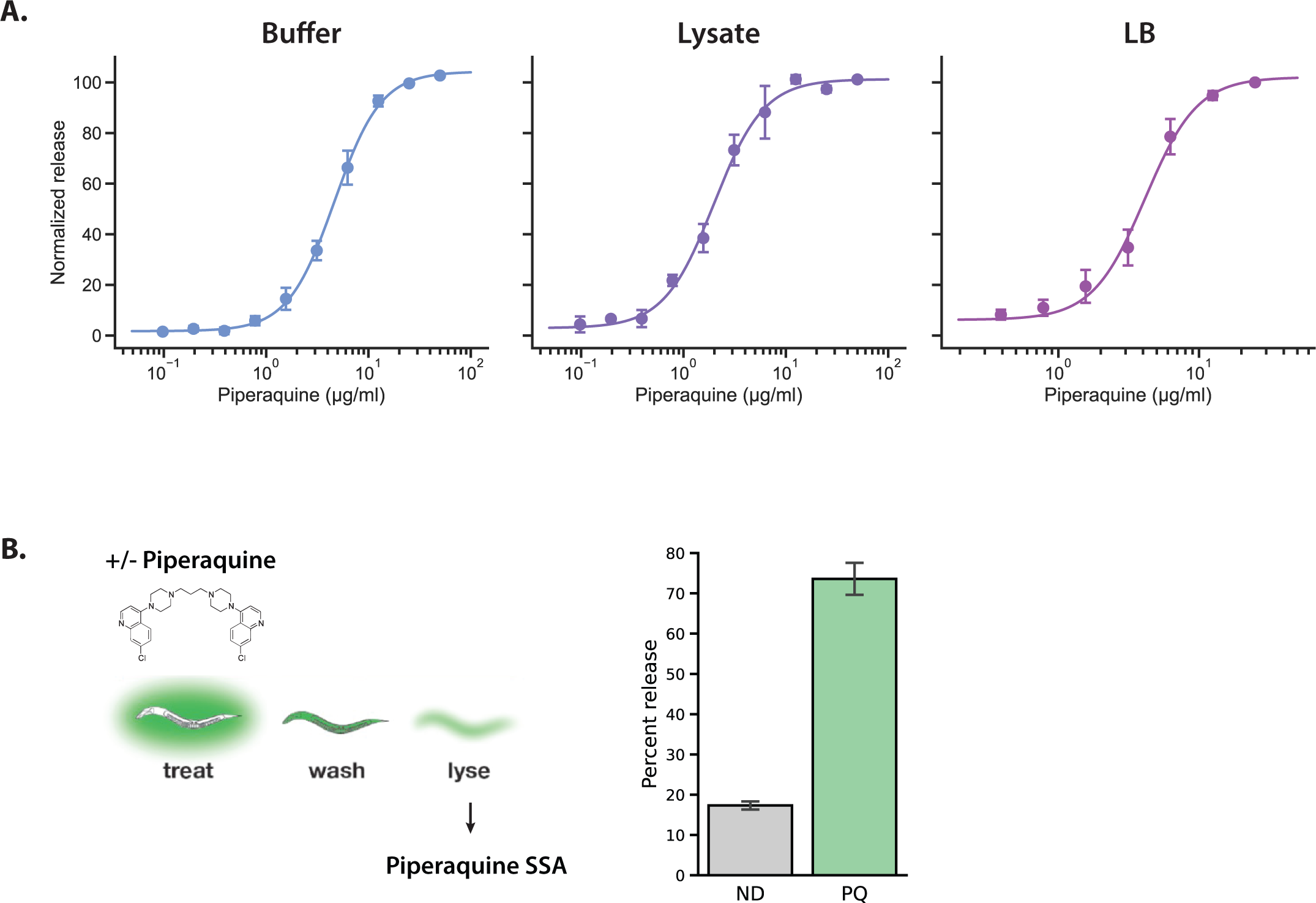
bSSAs work in cell lysate and other complex mixtures. **(A) Dose response of the piperaquine (PQ) bSSA in response to exogenous PQ added in buffer or other complex solutions.** Various amounts of PQ were titrated into either PBS buffer (left panel), worm lysate diluted 1:5 in PBS buffer (middle panel) or LB broth diluted 1:2 in PBS buffer (right panel). Mg^2+^ was added to all three conditions to a final concentration of 10 mM added Mg. Worm lysate was derived from N2 *C. elegans* worm pellet lysed in 3x volume of extraction solvent and diluted into PBS buffer. Dose response was plotted after subtracting the percent release at each PQ concentration from the percent release in either lysate or LB alone and normalized as further detailed in Methods. (B) Use of sensors to detect drug uptake in whole animal. *bus-5(br19) C. elegans* worms were treated with 100 μg/ml of piperaquine (in 5% methanol) for 7h in liquid culture (‘PQ’ sample). Worms treated with 5% methanol was used as the negative control (no drug ‘ND’ sample). After drug treatment, worms were washed with buffer, and the pellet lysed in 3x volume of extraction solvent. The worm lysate was then diluted 1 in 5 in PBS buffer, incubated with the PQ bSSA and the percent SRO release was plotted. Error bars represent standard error from 3 independent replicates.

### barcode SSAs can be multiplexed and act as independent sensors

bSSAs each release a different barcode and thus they should be able to be multiplexed to allow the parallel readout of many ligands simultaneously. To test this, we wanted to test different bSSA mixtures. In each case one bSSA can detect one ligand and releases the T7 SRO — this can be detected by fluorescence using our reporter assay (Fig 2). The other bSSA releases a different SRO that does not contain the T7 promoter and thus cannot drive fluorescence. We can thus use fluorescence to distinguish between the two bSSAs.

We tested multiplexing using 2 bSSAs — one detects piperaquine (PQ) and the other detects mefloquine (MQ) — that individually recognize different ligands (Fig 7A). We made two pools of these PQ and MQ bSSAs. In pool 1, the PQ bSSA is paired with a T7 SRO and the MQ with a T3 SRO; in pool 2 the MQ is paired with the T7 SRO and the PQ with the T3. Since the fluorescent readout is specific for T7 SRO release, pool 1 should only show signal if PQ is added and pool 2 only when MQ is added. That is exactly what we observe: addition of PQ resulted in fluorescence in pool 1, whereas addition of MQ resulted in fluorescence only in pool 2. This shows that bSSAs can be multiplexed and each responds independently to their ligand.

**Fig 7.**
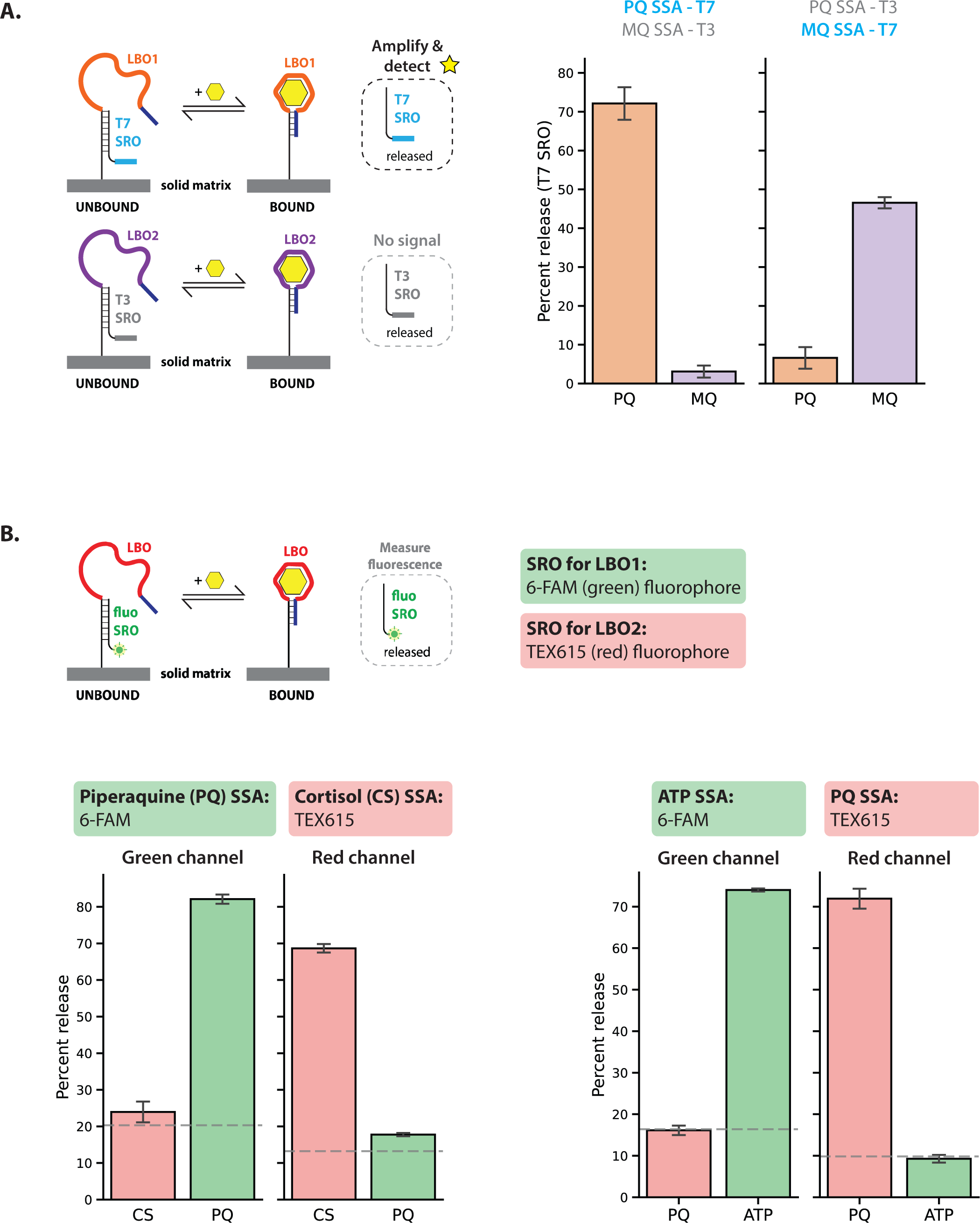
Ligand binding in a mixture of bSSAs. **(A) Specific barcode release in a mix of bSSAs that detect different ligands.** (Left) The piperaquine (PQ) LBO is paired to a T7 barcode SRO while the mefloquine (MQ) LBO is paired to a T3 barcode SRO. (Right) The PQ LBO is paired to a T3 barcode SRO while the MQ LBO is paired to a T7 barcode SRO. The two bSSAs were mixed in equal amounts then incubated with either 50 µM PQ or MQ. **(B) Multiplexing of bSSAs using different fluorophores.** A fluorescence readout was used where target binding induces a conformational change that results in displacement of a short oligo attached to a fluorophore. The fluorescence intensity of the released fraction is then directly measured to determine the amount of SRO released. (Left) A piperaquine (PQ)-binding LBO is paired with a 6-FAM SRO, while a cortisol-binding LBO is paired with a TEX615 SRO. The two SSAs were mixed in equal amounts then incubated with either 50 μM of PQ or 200 μM of cortisol in 5% ethanol (in PBS). (Right) An ATP-binding LBO is paired with a 6-FAM SRO, while a PQ-binding LBO is paired with a TEX615 SRO. The two SSAs were mixed in equal amounts then incubated with either 50 μM of PQ or 125 μM of ATP. Fluorescence intensity was then measured in each sample at 2 different excitation/emission wavelengths for 6-FAM (‘green channel’; ex 485 nm, em 520 nm) and TEX615 (‘red channel’; ex 584 nm, em 620 nm). Grey dotted lines indicate SRO release from buffer alone. Error bars represent standard error from 3 independent replicates.

Since these experiments only detect the signal from one of two bSSAs in each pool, we also tested multiplexing with fluorophores with non-overlapping spectra to measure the release of barcodes from two bSSAs simultaneously (Fig 7B). Here, we first pair a cortisol (CS) bSSA with an SRO attached to a TEX615 fluorophore and pair a PQ bSSA with an SRO attached to a 6-FAM fluorophore. Release of the TEX615 SRO can then be detected in the red channel and release of the 6-FAM SRO in the green channel. This allows us to monitor release of SROs from both bSSAs at the same time. As before, we mix both bSSAs and add either ligand to the pooled sample.

We show that piperaquine addition results in more 6-FAM SRO release while cortisol addition results in more TEX615 SRO release, again suggesting that each bSSA functions independently and responds specifically to their cognate ligand even when mixed with other bSSAs. We also carried out the same multiplexing experiment with a PQ bSSA paired with a TEX615 SRO and an ATP bSSA paired with a 6-FAM SRO and show again that each bSSA functions as an independent sensor and releases their SRO only in the presence of their cognate ligand (Fig 7B).

We conclude that it is possible to multiplex bSSAs and that each bSSA acts as an independent sensor in a pool of bSSAs. This opens the door to highly multiplexed libraries of bSSAs that can be used to detect many different metabolites or drugs in parallel.

### Barcoding structure-switching aptamers removes constraints on nucleic acid chemistry in the ligand recognition region

In this study we show that structure-switching aptamers can use barcode release as a readout for ligand binding — sequencing the released DNA barcodes reports on which ligands bound their SSA sensors. At first sight this looks unnecessarily complicated — if what you care about is which ligand-binding region detected its target ligand why not sequence the ligand-binding region? Why bother with the barcodes at all? If the LBO with the ligand binding region was purely made of DNA or RNA that is indeed true — the LBO has all the information needed.

However, nucleic acid chemistry is far more diverse than the naturally occurring bases we see in biology and includes XNAs (xeno nucleic acids) like HNA, FNA, LNA and many other derivatives^39^. Ligand binding by SSAs can be greatly improved by using these more diverse XNAs^40, 41^ but these cannot be sequenced directly and indeed reading XNAs requires specifically engineered polymerases^42^. Furthermore, while a polymerase can be engineered to be able to read one specific XNA^42^, there is no ‘universal’ polymerase that can read or sequence all XNAs or any combination of different XNAs. This greatly restricts the chemical space of the nucleic acids used in the ligand-binding region of the SSA.

A great advantage of using DNA barcodes to read out ligand binding to SSAs is that it frees the ligand-binding region to be able to include any XNA or indeed any combination of XNAs — as long as the barcode released from every bSSA is DNA, it provides a single chemistry for readout without restricting the chemistry for detection. To test that this works, we constructed a bSSA for PQ in which the common or ligand-binding region of the LBO uses a mixture of DNA and 2′-OMe-RNA bases (Fig 8A-B). We show that this mixed nucleic acid ligand-binding region responds to PQ by releasing its barcode (Fig 8B). The barcode SSA technology thus allows ligand recognition to involve any nucleic acid, whether naturally occurring or artificial. Since every bSSA is read out by release of DNA barcodes, bSSA libraries can include sensors with any possible nucleic acid in any possible combination within the ligand-binding region, but these can all be read out with the same basic DNA barcode chemistry. This provides immense flexibility and widens the nucleic acid space that can be used for ligand detection. We also note that it is also possible to attach the barcode to the LBO itself and either transcribe or release it as shown (Supp Fig 1A) and show that these architectures also give functional sensors (Supp Fig 1B-C). The principle is nonetheless the same: the nucleic acids in the ligand-binding region are not constrained and the readout of SSA ligand binding is released DNA barcodes.

**Fig 8.**
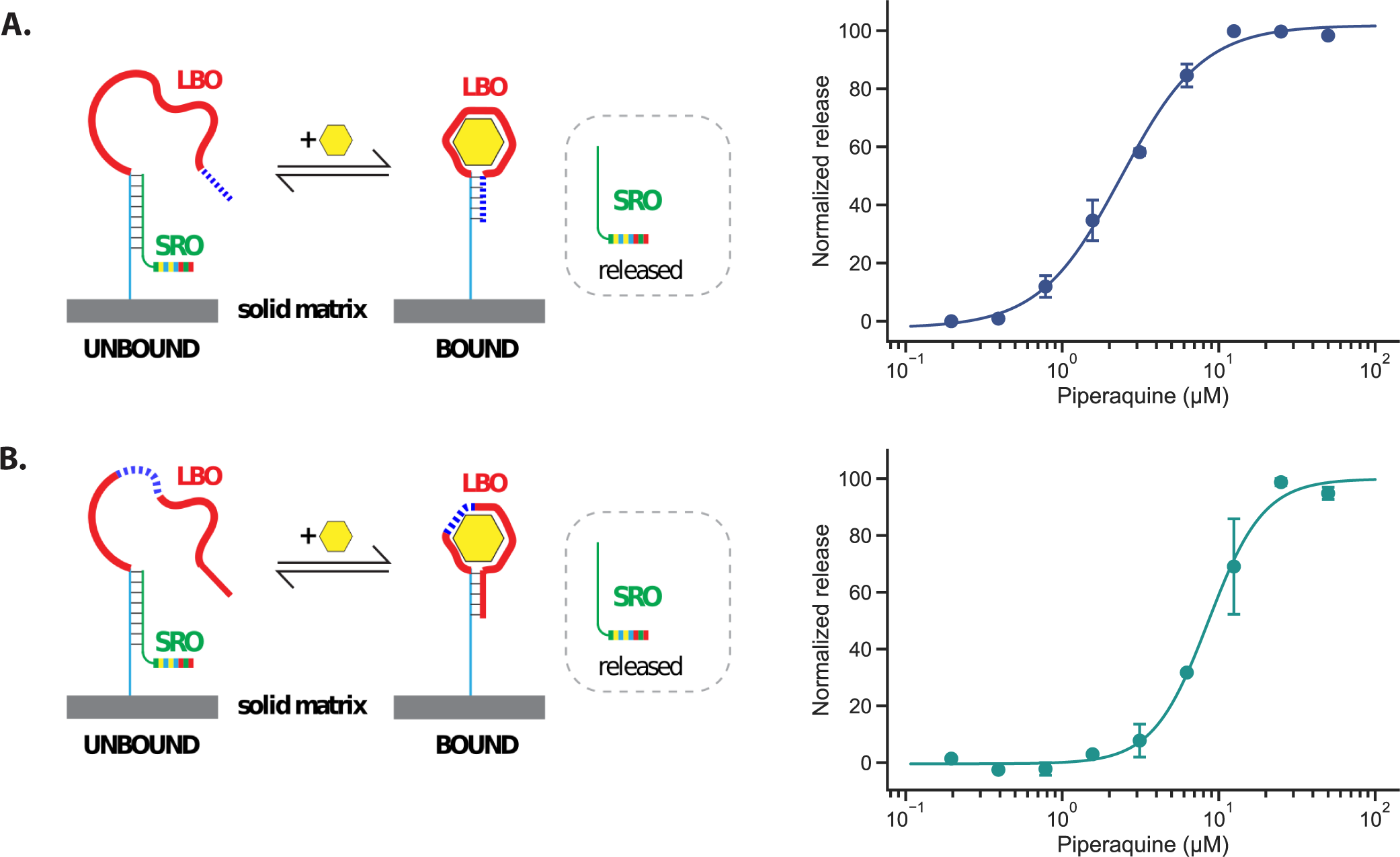
2ʹ-OMe-RNA/DNA chimeras can function as LBOs. **(A) PQ LBOs with DNA bases replaced by 2ʹ-OMe-RNA bases in the common stem region.** 8 nucleotides at the 3′ end of the LBO were replaced by 2ʹ-OMe-RNA bases (blue dotted region), as shown by the schematic on the left. Ligand binding was assayed as described previously and the dose response curve plotted on the right. **(B) PQ LBOs with DNA bases substituted by 2ʹ-OMe-RNA bases in the ligand-binding region.** 4 nucleotides in the ligand-binding region (blue dotted region) of the LBO were replaced by 2ʹ-OMe-RNA bases (blue dotted region), as shown by the schematic on the left. These bases are predicted by mFold^47^ to be base-paired and were chosen to minimise the likelihood of disrupting specific ligand contacts. The corresponding dose response is plotted on the right. Error bars represent standard error.

In summary, we believe we have established a new platform for multiplexed detection of metabolites and drugs using structure-switching aptamers with a DNA barcode readout. The released barcodes can be highly amplified allowing signal amplification to improve detection of low abundance targets. Since the readout for all sensors is DNA barcode release, the system is fundamentally compatible with any platform for single-cell or spatial transcriptomics and thus opens a new dimension for single cell multiomics. Finally, unlike many SSA configurations, this allows the use of any single XNA or any combination of XNAs in the ligand-binding region while still reading out ligand-binding with DNA barcode release. It is thus possible to make large collections of barcode SSAs each with a very different nucleic acid chemistry in their ligand-binding regions and read all out in parallel on a single platform.

## Discussion

Single-cell genomics has been a major advance. Single-cell RNA-seq has transformed our view of development^43, 44^ and single-cell genome sequencing has shed new light on tumorigenesis^45^. Ultimately, single-cell genomics relies on the power of PCR to amplify the limited material available in a single cell. For metabolites and drugs, however, single-cell approaches are far more challenging since there is no current way to amplify these small organic molecules. While mass spectroscopy is now very sensitive, it remains extremely challenging to do single-cell metabolomics. Here we establish a new platform with power to transform metabolomics to the scale of single cells: barcode structure-switching aptamers (bSSAs).

Each bSSA recognises a specific ligand and is coupled to a specific DNA barcode. Ligand binding drives a conformational change that releases the barcode. By sequencing released barcodes, we can determine which bSSAs detected their cognate ligands and thus we can map the complex molecular space of metabolites and drugs into the simple sequence space of DNA barcodes. These can be amplified using routine methods used in most other single-cell genomics — thus it is possible to detect and quantify DNA, RNA, and metabolites on a single platform.

In this study, we show that we can use bSSAs to specifically measure levels of different classes of small molecules, and that novel or existing aptamers can be easily adapted to be barcode SSAs using the architecture we describe here. We show that each bSSA behaves as an independent sensor in a mixture of bSSAs — this makes it possible to have multiplexed detection of many hundreds of ligands in the same assay volume since each releases a different barcode. Crucially, we show that these bSSAs can work even in the context of extremely complex mixtures of biomolecules such as cell lysates. Finally, we report methods to generate barcode SSAs which allows the correct matching of ligand-binding oligo and barcode release oligo in bulk. This allows the efficient generation of large numbers of independent barcode SSA sensors without having to assemble them individually.

Barcode SSAs thus allow the multiplexed detection of many different metabolite and drug ligands in complex biological mixtures such as cell lysates and PCR can be used to amplify the barcodes and greatly increase the power of detection. We believe that barcode SSAs will allow DNA, RNA, and metabolites to be read on a single platform at the scale of single cells.

## Acknowledgements

This research was funded by project grant PJT183712 to A.G.F. from the Canadian Institute of Health Research (CIHR). We also thank members of the Fraser Lab for advice and helpful discussions throughout. *C. elegans* strains were provided by the CGC, which is funded by NIH Office of Research Infrastructure Programs (P40 OD010440).

## Competing Interests

The authors declare the following competing interests: A.G.F., J.H.T. and M.P.M. have patents pending related to the use of barcode structure-switching aptamers.

## Materials and Methods

### Reagents

All oligonucleotides were synthesized by Integrated DNA Technologies (IDT) (Coralville, IA) and dissolved in nuclease-free water at a concentration of 100 µM. The oligonucleotide sequences used were adapted from Yang *et al.*^20, 35^; Coonahan *et al.*^38^; Song *et al.*^37^; Warner *et al.*^31^; and Huizenga & Szostak^36^. Oligonucleotide sequences used for bSSAs are listed in Supplemental File 1. Stock solutions of amino acids, antibiotics, ATP and pentamethylcyclopentadienyl rhodium dichloride dimer [Cp*RhCl_2_]_2_ (Cp*Rh(III), 5 mM) were made in nuclease-free water while the stock solution of tyrosine (2 mM) was made directly in binding buffer (20 mM HEPES (pH 7.5), 1 M NaCl, 10 mM MgCl_2_, 5 mM KCl). Stock solutions of piperaquine (1 mg/ml) and mefloquine (500 μg/ml) were made in 5% methanol. Stock solutions of cortisol were made in ethanol with a final working concentration of 5% ethanol. The stock solution of DFHBI-1T (20 mM) was made in DMSO.

### Ligand binding assay

For each sample, 5 µL of Dynabeads MyOne Streptavidin C1 magnetic beads (Invitrogen) were washed 3x in Bind & Wash buffer as per manufacturer’s protocol, and finally resuspended in 10 μL binding buffer. 25 pmol of LBO and 125 pmol of SRO (10 μL total, diluted in binding buffer) were heated at 95°C for 5 min and slowly cooled to 25°C. The oligos were then added to the resuspended beads and incubated on a rotator for 30 min at room temperature. The beads were then washed 2x with 100 μL binding buffer and resuspended in 20 μL of the same buffer and incubated for another 45 min. The beads were then washed 1x and resuspended with various concentrations of ligand. The samples were incubated for 45 min on a rotator at room temperature and the supernatant was collected (‘supernatant’ sample). The beads were then resuspended in 20 μL of strand separation buffer (20 mM HEPES (pH 7.5), 300 mM NaCl), heated at 95°C for 4 min, and the supernatant sample collected (‘remainder’ sample). Percent release is the calculated as: 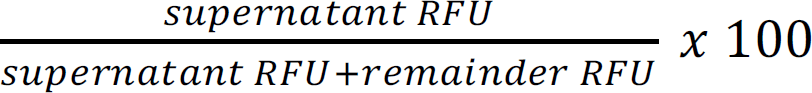.

For experiments with phenylalanine, tyrosine, leucine and tryptophan, the amino acids were first complexed with Cp*Rh(III) at a final concentration of 100 μM Cp*Rh(III) and varying concentrations of amino acids as described by Yang *et al.* ^20^, and incubated together at room temperature for >45 min. For experiments with piperaquine and mefloquine, the beads and oligonucleotides were prepared as described above, except that PBS with 10 mM MgCl_2_ was used as the binding buffer, and PBS was used as the strand separation buffer.

For the LB experiments, the LB broth (BioShop) was diluted 1:2 in PBS + 10 mM Mg and spiked with various amounts of piperaquine. SSAs were then incubated in these mixtures. LB medium with no piperaquine added was used as the negative control. For lysate experiments, lysates of *C. elegans* worms were diluted 1:5 in PBS + 10 mM Mg and spiked with various amounts of piperaquine. Lysate with no piperaquine added was used as the negative control.

### SRO readout

For samples with a T7 SRO, 1.5 µL of the 20 µL supernatant sample was added to 5 µL of nuclease-free water and 2 µL of 10 µM Baby Spinach transcription template. Oligos were mixed and heated at 95°C for 5 min and slowly cooled to 25°C. Reaction buffer, NTPs (2 mM each) and T7 RNA polymerase were then added to the sample for a total volume of 20 µL and incubated at 37°C for 2h. After transcription, the sample was then heated at 90°C for 2 min before incubation on ice for >3 min. 30 µL of water and 5 µL of 100 µM DFHBI-1T (diluted in Tris-HCl buffer: 40 mM Tris-HCl (pH 8.0), 5 mM MgCl_2_, 125 mM KCl) were added to 15 µL of the sample, and the sample was then heated to 65°C for 5 min and slowly cooled to room temperature (as per Okuda *et al.*^33^). Each sample was then transferred to a 96-well plate and the fluorescence intensity for each sample was measured using a FLUOstar Omega microplate reader (excitation 485 nm, emission 520 nm).

For samples with a 6-FAM or TEX615 SRO, 20 µL of the sample was added to 35 µL of water and fluorescence was directed measured using the microplate reader (6-FAM: ex 485 nm, em 520 nm; TEX615: ex 584 nm, em 620 nm).

### Two-color multiplexing assay

The piperaquine LBO was paired with an SRO attached to a 6-FAM fluorophore (IDT) while the cortisol LBO was paired with an SRO attached to a TEX615 fluorophore (IDT). For the multiplexing experiment involving piperaquine and cortisol, two different stem variants were used for the piperaquine and cortisol SSAs, and equal amounts of piperaquine and cortisol SSAs bound to beads were mixed together. Ligand binding was then assayed in PBS supplemented with 10 mM Mg and 5% ethanol. SSAs were incubated with either piperaquine or cortisol for 45 min on a rotator at room temperature and the supernatant was collected. The beads were then resuspended in 20 µL of PBS, heated at 95°C for 4 min, and the supernatant collected. After addition of 35 μl water to each sample, fluorescence was then measured using the microplate reader at 2 different excitation/emission wavelengths for 6-FAM (ex 485 nm, em 520 nm) and TEX615 (ex 584 nm, em 620 nm).

### Drug treatment and worm lysis

*C. elegans* worms (wild-type N2 strain and *bus-5(br19)*, which shows increased drug permeability^46^) were grown and maintained on NGM agar plates seeded with OP50 bacteria. Strains were provided by the CGC, which is funded by NIH Office of Research Infrastructure Programs (P40 OD010440).

For drug treatments, OP50 cultures were heat-killed at 65°C for 30 min, spun down, then concentrated 2-fold in NGM media. 1600 μl of the suspended cultures was then added to 200 μl of mixed stage *bus-5(br19)* worms in M9, along with 200 μl of 10x concentrated drug. The worms were then incubated with the drugs for 7h in a 20°C shaker, with piperaquine and mefloquine added at a final concentration of 100 μg/ml and 50 μg/ml respectively. Worms were treated with 5% methanol in lieu of drugs as the negative control. After 7h drug treatment, the worms were washed 3x in M9 followed by 1x in PBS. The worm pellet was then flash frozen and stored at -80°C.

To lyse the worms, the frozen pellets were ground with a pestle until the pellet defrosted. Extraction solvent (8:1:1 ratio of methanol:chloroform:water) was then added to the tubes at 3x the volume of the pellet. Samples were then vortexed and subjected to 3 freeze-thaw cycles. The tubes were then centrifuged at 13,200 rpm for 15 min and the supernatants were collected and stored at -80°C until needed. The worm lysate used for the piperaquine spike-in experiment was collected from N2 worms that were not treated with any drugs.

### Cleavage of abasic site

A dSpacer (IDT) was used as the abasic site. For each sample, 25 pmol of SSAs were folded in 1x NEBuffer 3 (NEB) by heating at 95°C for 5 min followed by slow cooling to 25°C. Endonuclease IV (NEB) was then added to the sample and the reaction mix was incubated at 37°C overnight. After incubation, the sample was then added to the dynabeads (described above), along with additional NaCl and KCl such that their final concentrations were at 1 M and 5 mM respectively. The binding assay was then carried out the same way as above.

---

**Supp Fig 1.**
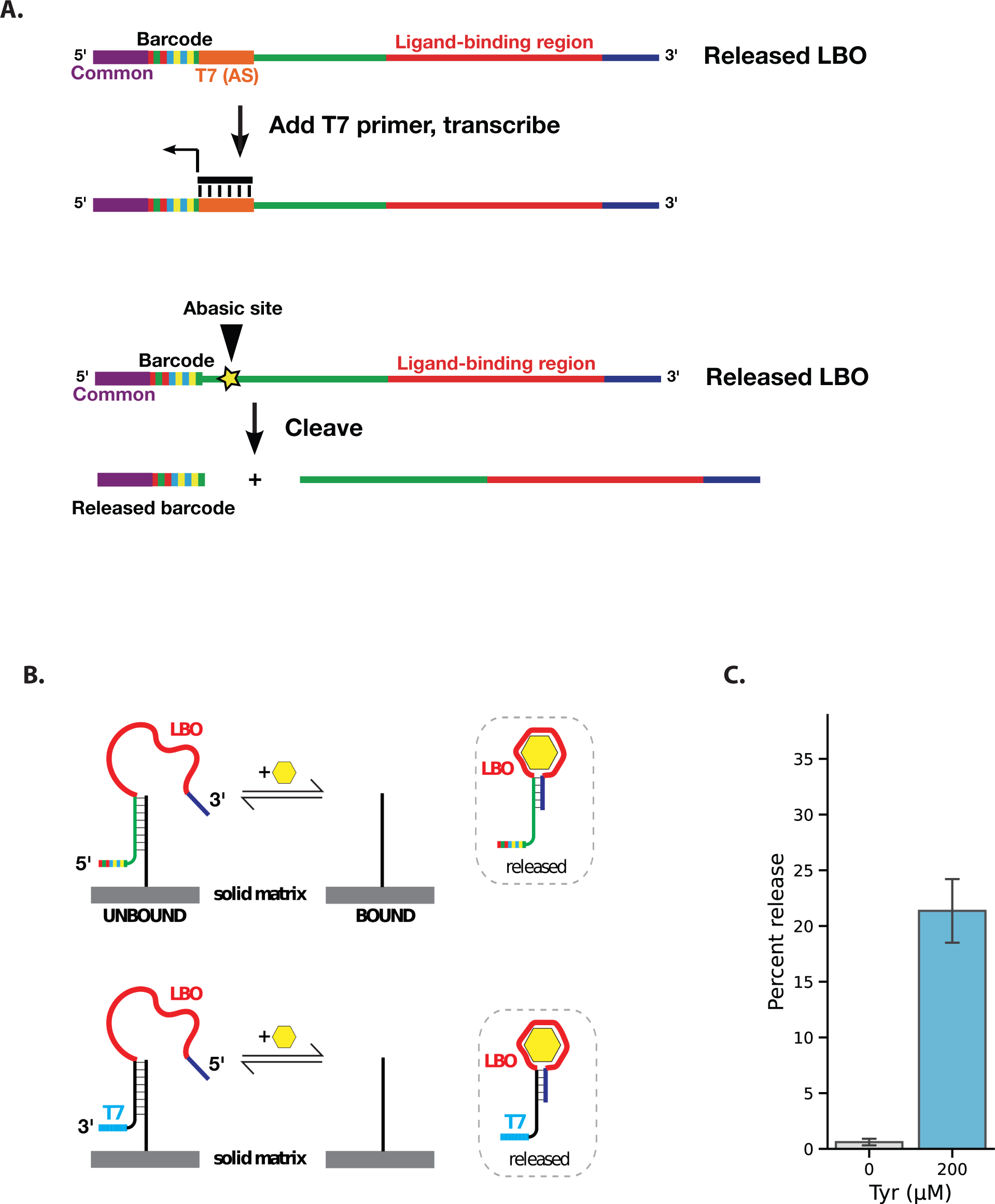
Barcode SSAs can also be configured with an immobilized short common oligo and a released LBO. **(A) Alternative methods to separate barcode from the rest of the LBO.** Antisense versions of the barcode and T7 promoter can be placed upstream of the ligand-binding region of the LBO. T7 primers could then be added to transcribe and amplify the barcode region. In this way, only the regions consisting of DNA bases need to be read. An abasic site or restriction site could also be cleaved to separate the barcodes from the rest of the LBO. (B) Re-configuration of the T7-Baby Spinach assay to assay release of LBOs in this alternative configuration. (Top) Barcodes can be added to the 5ʹ end of LBOs so that they can be released and transcribed as shown in (A). (Bottom) The 5ʹ and 3’ common ends of the LBO were flipped so that the T7 promoter is present at the 3’ end of the LBO. The LBO sequences released upon ligand binding were then used to transcribe a Baby Spinach template as described above. (C) Release of a Tyr T7 LBO upon ligand binding. A tyrosine T7 LBO is released and detected in our assay upon addition of 200 μM tyrosine. Error bars represent standard error from 7 independent replicates.

## Notes

### Competing Interest Statement

A.G.F., J.H.T. and M.P.M. have patents pending related to the use of barcoded structure-switching aptamers.

